# Local negative frequency-dependence can decrease global coexistence in fragmented populations

**DOI:** 10.1101/2025.05.19.654891

**Authors:** Anush Devadhasan, Oana Carja

## Abstract

Most biological populations are rich in diversity, and negative frequency-dependent (NFD) selection is a well-established mechanism thought to underlie this stable coexistence of multiple variants. Recent studies confirm its widespread presence at local spatial scales, however it remains unclear whether these local-scale dynamics are sufficient to maintain biodiversity across larger, landscape-level scales. While prior theoretical work has found that local NFD selection can indeed promote global coexistence, these studies only analyzed contiguous landscapes. In contrast, many ecosystems are not contiguous, but rather spatially fragmented or exhibit spatial variation in the local carrying capacity. Using a theoretical model based on the classic island framework, we show that in fragmented populations, NFD selection can paradoxically reduce coexistence and shorten fixation times, relative to neutrality. Fragmentation also produces a non-monotonic relationship between fixation time and population size, with diversity lowest at intermediate scales, in contrast to the classical species-area relationship. We show that these results persist in a multispecies modeling framework. We also develop a statistical test to detect whether NFD selection suppresses coexistence in fragmented systems, and apply it to a presence-absence dataset of avian species in the Ryukyu Islands, finding evidence that NFD selection indeed reduces biodiversity in this case. Together, our findings suggest that fragmentation can undermine the stabilizing effects of NFD selection, calling into question its generality as a mechanism for maintaining biodiversity in heterogeneous landscapes.

**Broader significance:** As human activities increasingly fragment once-continuous habitats, it becomes ever more critical to understand how these changes impact biodiversity. Negative frequency-dependent (NFD) selection, a key mechanism promoting coexistence, is well known for preserving genetic and species diversity in well-mixed and spatially continuous populations. Using a theoretical model of a subdivided population, we show that NFD selection can actually accelerate extinctions and erode biodiversity, if the population is fragmented. This counterintuitive result arises because habitat fragmentation disrupts the alignment between the local scale of ecological interactions and the broader landscape structure of the habitat. When local frequency-dependent dynamics no longer scale up across the landscape, their stabilizing effect is lost. Our findings suggest that, in fragmented environments, biodiversity may be better preserved under neutral dynamics or even weak directional selection, than under NFD selection.

## Introduction

The maintenance of biodiversity across the variety of natural systems has been a topic of intense fascination in ecology and evolutionary theory alike. From genes to species to entire ecosystems, biological systems are replete with variants that manage to stably coexist. Over the years, numerous mechanisms have been proposed to explain how such diversity is maintained in the face of selection, competition, and limited ecological resources, culminating in the development of Modern Coexistence Theory (MCT) (Chesson, 2000). MCT provides a unifying framework that distills the problem of coexistence into the balance between two key forces: (1) niche differentiation, which promotes coexistence by reducing interspecific competition relative to intraspecific competition, and (2) fitness differences, which diminish coexistence by promoting competitive exclusion (Chesson, 2000; Barabás and D’Andrea, 2020). Empirical studies across a wide range of ecosystems have subsequently demonstrated the applicability and predictive power of MCT in explaining observed patterns of coexistence (Letten et al., 2021; Stanley Harpole and Tilman, 2006; Anderson et al., 1981). At a big picture level, niche differentiation in MCT is able to maintain coexistence because it gives rise to negative frequency-dependent (NFD) selection, where the fitness of a species or genetic variant decreases as it becomes more common (Vellend, 2010). This form of balancing selection is an important mechanism of coexistence across biological scales (Christie and McNickle, 2023; Takahata and Nei, 1990; Huang et al., 2012), with ample empirical support (Radwan et al., 2020; Kurbalija Novičić et al., 2020; Fitzpatrick et al., 2007).

Separately, building on the foundational ideas described by the neutral theory of molecular evolution, another minimal model that can explain patterns of coexistence is neutral theory. In this framework, diverse communities of equivalent species emerge when random extinctions are balanced by speciation or mutation events (Hubbell, 2005; Leibold et al., 2004). Unlike the deterministic, infinite population size models frequently used to study NFD selection, where coexistence can be maintained indefinitely, in these finite population neutral models, genetic drift inevitably leads to the fixation of a single variant, in the long term, and one proxy for coexistence is the conditional fixation (or equivalently persistence) time (Schreiber, 2017; Schreiber et al., 2023). Direct comparison of these diversity-promoting mechanisms has shown that various indicators of coexistence and biodiversity (including fixation times) are significantly higher under NFD selection compared to neutrality (Pigolotti and Cencini, 2013; Carroll and Nisbet, 2015; Huang et al., 2012). However, for populations with spatial structure, the ecological processes driving NFD selection, such as Janzen-Connell effects, occur over local spatial scales and are governed by local frequencies, with dynamics that might not always scale up to ensure coexistence at larger, global levels (Hülsmann et al., 2021). For instance, a species might be rare, and gain a fitness advantage, within a local patch, even if it is globally abundant (Molofsky et al., 1999). Therefore, although local NFD selection has been documented across a range of ecosystems, including tropical and temperate forests, grasslands, coral reefs, microbial communities, and vertebrate populations (Kalyuzhny et al., 2023; LaManna et al., 2017; Bagchi et al., 2014; Johnson et al., 2012; Maron et al., 2016; Marhaver et al., 2013; Dimitriu et al., 2019; Madsen et al., 2022), such evidence alone does not imply that local NFD dynamics maintain global diversity and theoretical models are needed to bridge this gap.

While, traditionally, theoretical frameworks used to study coexistence and NFD selection have been developed independently from the frameworks used to study the effects of spatial structure, such as metacommunity theory or evolutionary graph theory (Kuo and Carja, 2024), recent work has begun to investigate the role of spatial structure in shaping patterns of biodiversity that can be maintained by NFD selection by analyzing models incorporating contiguous, regular spatial landscapes. At a big picture level, these studies find that local NFD selection does, in fact, continue to maintain high levels of global coexistence (May et al., 2020). However, these studies also show that when small, interspecific differences exist, they can quickly overwhelm the stabilizing effects of frequency-dependence and dramatically reduce coexistence. For example, discrete lattice and continuous planar Lotka-Volterra models have found that spatial structure shrinks the coexistence region of the parameter space, even when the classical condition, intraspecific competition larger than interspecific competition, holds (Neuhauser and Pacala, 1999; Murrell, 2010). Stochastic multispecies models in continuous planar (May et al., 2020) and discrete lattice-structured (Miranda et al., 2015; Stump and Comita, 2018) spatial representations have further shown that interspecific differences in the strength of NFD selection can sharply reduce biodiversity proxies like fixation times, species richness, and evenness. Similarly, lattice-based models have demonstrated that fitness asymmetries (Chisholm and Fung, 2020; Smith, 2022) and differences in resource uptake rates (Weiner et al., 2019) can also significantly diminish diversity relative to well-mixed environments.

This prior work, however, has so far only focused on contiguous, regular spatial landscapes. In contrast, many natural ecosystems are fragmented or patchy, due to habitat loss (Haddad et al., 2015) or spatial heterogeneity in local carrying capacity (M’Gonigle et al., 2012). This fragmentation is likely to increase the disconnect between local interactions and global dynamics, potentially weakening the stabilizing potential of NFD selection even further. Although fixation-time studies have examined fragmented systems (e.g., island or network-structured populations) (Slatkin, 1981; Cherry and Wakeley, 2003; Hindersin et al., 2016; Kuo and Carja, 2024), most focus on neutral or constant selection, with only limited work on frequency dependence in studies of cooperative games (Hauert et al., 2014; Ying et al., 2018). We lack a general theory for how NFD selection shapes patterns of coexistence in fragmented spatial systems and how properties of population spatial arrangement modulate the relative strength of NFD selection versus neutrality in shaping and maintaining patterns of coexistence.

Here, we introduce a stochastic model of local frequency-dependent selection in an island-structured population and ask how this population fragmentation shapes patterns of coexistence, compared to neutrality. Building on the classic island model, we relax the standard assumption of constant selection (Wright, 1931, 1940). To quantify coexistence, we first analyze a tractable two-variant model and study the average fixation time of a mutant variant invading a wild-type population using simulations and analytic approximations. We find that in fragmented populations, for small enough migration rates, fixation times (and thus coexistence) are shorter under NFD selection than under neutrality, reversing the trend observed in well-mixed systems. Moreover, in this regime, fixation times also show a non-monotonic dependence on deme size.

While our simple two-variant model facilitates analytic tractability, to generalize these findings, we extend our analysis to a multi-species model, using species richness as a proxy for coexistence. We show that, under the same parameter regime, NFD selection again leads to lower species richness than neutrality, consistent with analytical predictions from the two-variant case. These results enable us to develop a test for identifying whether natural populations operate in a regime where NFD selection reduces biodiversity, relative to neutral expectations, using only species presence-absence data. Applying this test to a dataset of avian species in the Ryukyu Islands (Matthews et al., 2023), we find evidence that NFD selection, and by implication, niche effects, is operating and shaping patterns of coexistence in this fragmented ecosystem.

Our results demonstrate that local NFD selection can decrease global biodiversity, relative to neutrality, under sufficient spatial fragmentation and that niche differentiation may be unable to maintain coexistence in fragmented habitats. This contrasts with prior models of well-mixed populations or models in contiguous space, where biodiversity losses under NFD selection arise only from interspecific differences, not from NFD selection itself. Our results may help explain why functional traits, which are indicative of niche differentiation, frequently fail to predict patterns of coexistence in natural communities (Kraft et al., 2015; Adler et al., 2013; Levine et al., 2025).

## Results

### A model of local frequency-dependent dynamics in fragmented populations

To understand the interaction between spatial structure and frequency-dependent selection, we first make use of an extension of Wright’s classic island model and consider a population composed of two different types of individuals: wild-type and mutant variants. The population is structured into D demes of N individuals each, with probability of migration m between any two demes. This means that, at every generation, an offspring’s parents come from within the deme with probability (1 – m) and from the overall population with probability m. The case when migration m = 1 represents a well-mixed population. We model neutral and frequency-dependent dynamics by assuming that reproduction within a deme is biased by local frequencydependent selection, such that the mutant fitness in deme i is given by 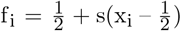, where x_i_ is the mutant frequency in deme i and s is a frequency-dependent selection coefficient (Molofsky and Bever, 2002). Notice that, if s is not zero, this mapping leads to a mutant equilibrium frequency of exactly 0.5 in the well-mixed population regime (see insert **Figure 1B**), although eventually drift will push the mutant to fixation or extinction. This flexible model, although simple, allows us to capture both structured (0 < m < 1) and well-mixed populations (m = 1), as well as neutral (s = 0), negative frequency-dependent (s < 0) and positive frequency-dependent (s > 0) selection (**Figure 1A**).

**Figure 1:**
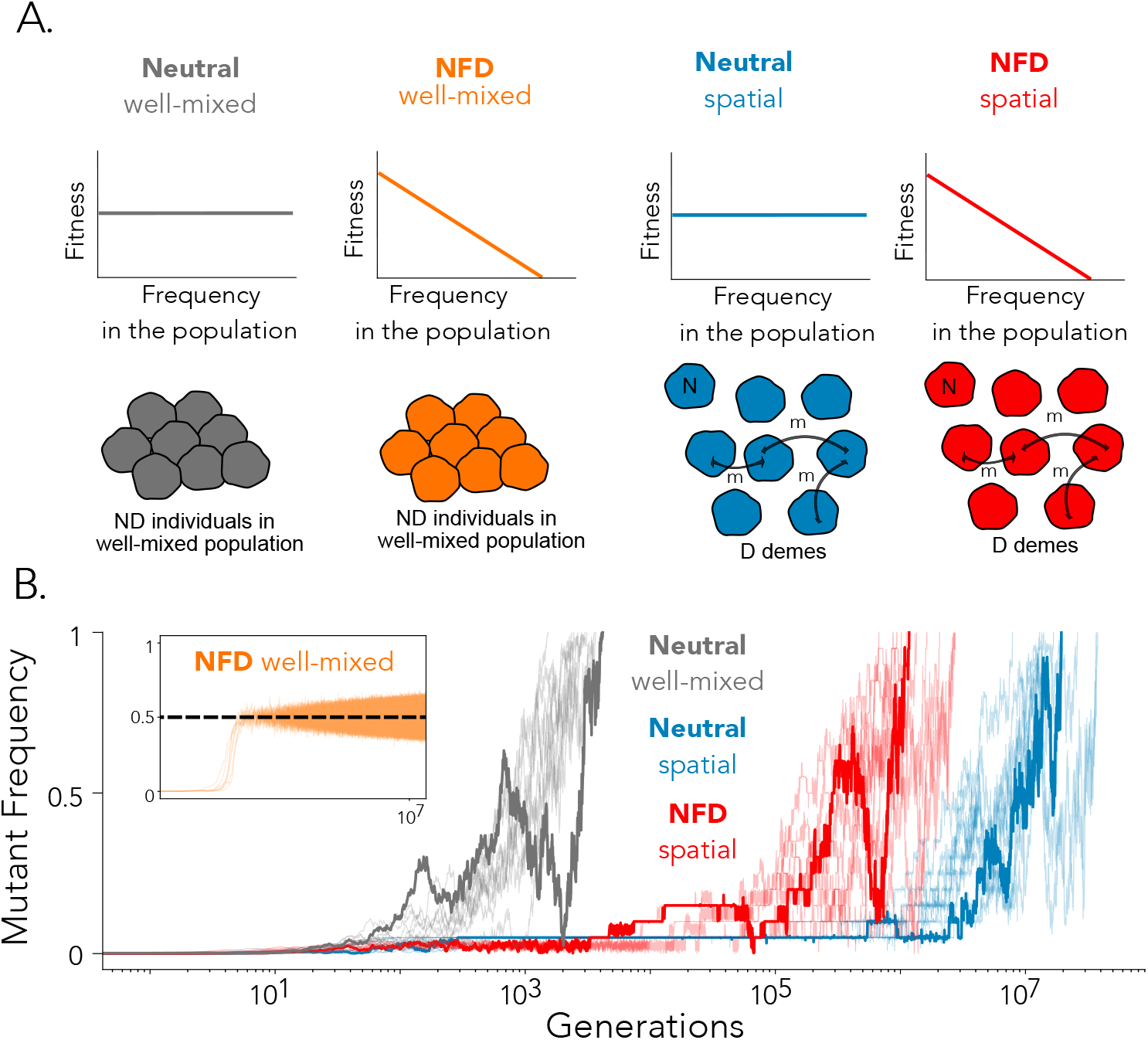
**Panel A**. Illustration of the four regimes we compare using our model. **Panel B**. Ten mutant trajectories for each of the four regimes considered. The total population size here is N_T_ = ND = 2000 with N = 100, D = 20 and m = 10^−6^ for the fragmented populations. Under neutrality, s = 0 and under NFD, s = –0.0625. One representative trajectory for each case is shown in bold.

Let us consider the average mutant fixation time (conditional on mutant fixation) as a proxy for coexistence in this model. While neutral dynamics are often invoked as a highly effective minimal model to explain patterns of coexistence (Hubbell, 2005; Leibold et al., 2004), the amount of biodiversity observed under neutrality, in the presence of genetic drift or demographic stochasticity, is much smaller compared to what can be observed under classic mechanisms of coexistence that give rise to negative frequency-dependent (NFD) selection, such as stabilizing niche differences. We visualize these phenomena in **Figure 1B**, where we simulate trajectories of a novel mutant variant in the population, under neutral (s = 0, m = 1) and negative frequency-dependent selection (s < 0, m = 1) in a well-mixed population (grey and orange trajectories, respectively). While under neutrality the mutant fixes after ∼ 10^3^ generations, there is still no fixation even after 10^7^ generations under negative frequency dependent selection.

In fragmented populations, even under neutral dynamics, spatial structure can significantly prolong coexistence compared to the well-mixed model, by restricting gene or species flow (grey (s = 0, m = 1) and blue (s = 0, 0 < m < 1) trajectories in **Figure 1B**) (Cherry and Wakeley, 2003). However, when both mechanisms are present (that is, in fragmented populations with local negative frequency-dependent selection, s < 0, 0 < m < 1) we find, counterintuitively, that the mutant tends to fix *faster* (∼ 10^6^ generations) than under either mechanism alone.

In **Figure 2** we study the amount of population fragmentation that can lead to this observed interplay effect. We first show the difference in fixation times between the frequency dependent and the neutral regimes for a well-mixed population. As expected, increasing the strength of NFD (negative values of s) increases fixation times, while increasing the strength of PFD (positive values of s) has the opposite effect, leading to a decrease in the fixation times (**Figure 2A**). As population fragmentation decreases, the result is maintained (**Figure 2B**), however, for small enough migration rates m (high enough levels of population fragmentation), the opposite result starts to be observed: negative frequency dependence decreases fixation times compared to neutrality, while positive frequency dependence increases fixation times and coexistence of the two variants in the population (**Figure 2C**).

**Figure 2:**
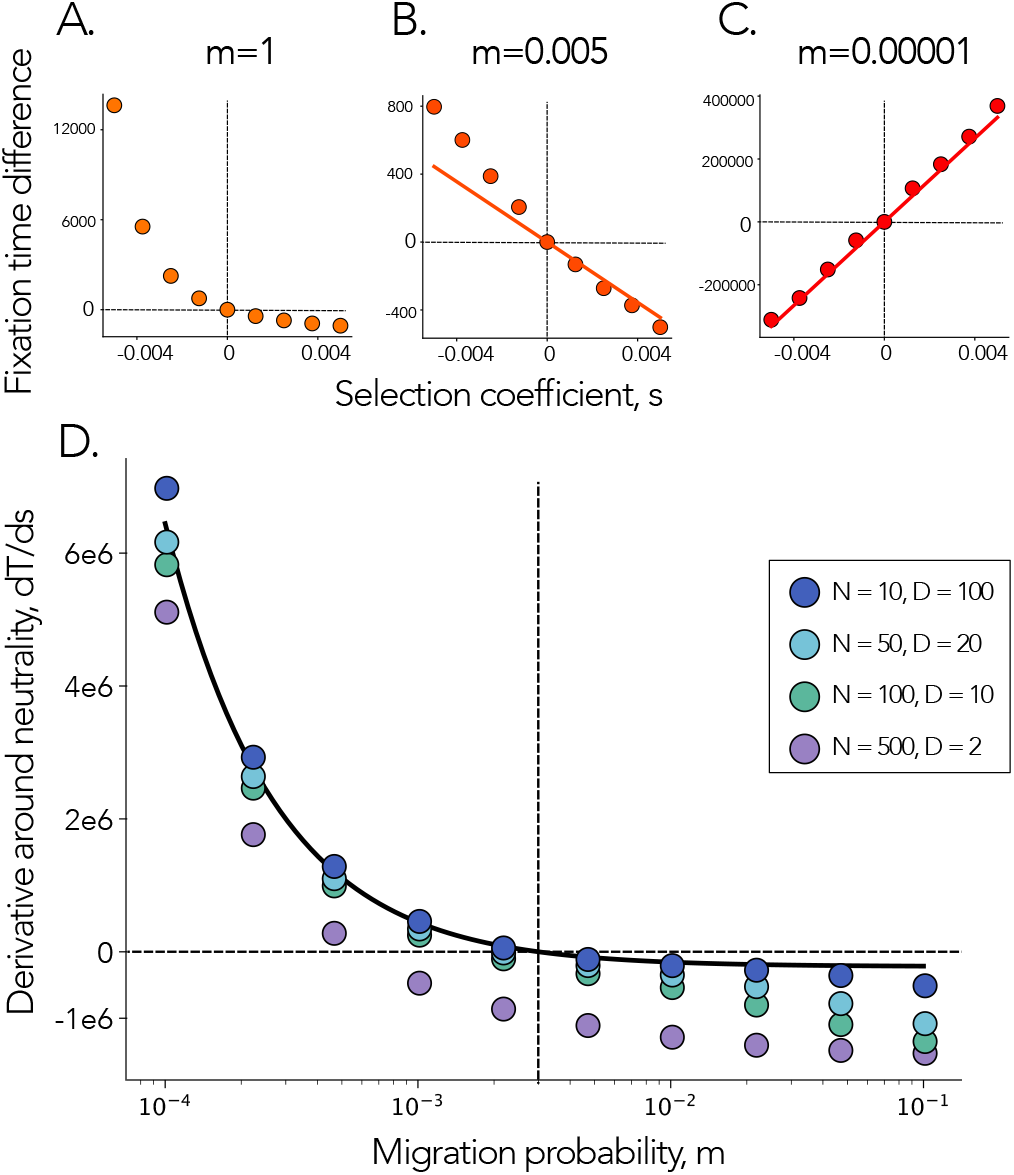
Fixation times and critical migration rates. We show the difference in fixation times relative to neutrality as a function of s in a well-mixed population in **Panel A**, and spatially-structured populations with m = 0.005 and m = 0.00001 in **Panels B** and **C**, respectively. Here N = 20 and D = 50. Dots represent averages over 10^6^ simulation runs, and solid lines show analytic approximations (equation (3)). **Panel D** shows the derivative of the fixation time with respect to the frequency-dependent selection coefficient s, as a function of migration rate m. Total population size is fixed to N_T_ = 1000, with deme size N and number of demes D varied as in the legend. Dots represent averages over 10^6^ simulation runs, while the black line represents equation (4).

To quantitatively understand this result, we derive analytic approximations for the conditional fixation time, assuming weak migration m << 1 (**Supplementary Appendix 1**). We first write analytic expressions for the mean per generation change in the mutant frequency, 𝔼(Δx),

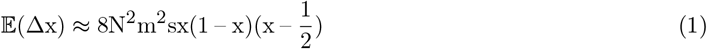

and the variance in the mutant frequency change per generation, Var(Δx),

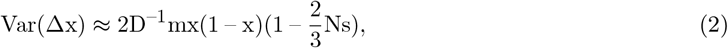

(see **Supplementary Appendix 1**). For s < 0 (the NFD regime), the sign of 𝔼(Δx) is determined by the sign of 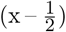 and increasing the strength of negative frequency dependence increases the magnitude of the selective force pushing the population towards the equilibrium frequency of 0.5, which effectively acts to increase the fixation time. In contrast, the expression for Var(Δx) shows that increasing the strength of NFD *increases* the strength of random fluctuations in the mutant frequency, which in turn acts to decrease the fixation time. More specifically, the effect of NFD on Var(Δx) comes from the 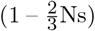 term, which becomes larger as s becomes more negative.

Since NFD simultaneously increases fixation time through 𝔼(Δx) and decreases fixation time through Var(Δx), the overall effect of NFD on the fixation time depends on the balance between these two effects. This balance is dictated by the strength of migration, m. Because 𝔼(Δx) scales with 𝒪 (m^2^) and Var(Δx) scales with 𝒪 (m), for large enough migration rate m, both the mean and variance in the per-generation mutant frequency change are of comparable magnitude. Under increasingly small migration however, 𝔼(Δx) vanishes faster than Var(Δx) and, since 𝔼(Δx) → 0, the population effectively evolves neutrally at a global spatial scale (although being under NFD over local spatial scales), while also experiencing stochastic fluctuations that are larger than what is observed under neutrality.

Using a diffusion approximation (Cherry and Wakeley, 2003), we write the average conditional fixation time of the mutant

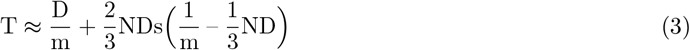

for small s << 1 and m << 1 (**Supplementary Appendix 1**) and show this approximation in **Figure 2B**, together with simulation results, in the large m regime. Although NFD still increases fixation time relative to neutrality here, we find that this increase is linear compared to the super-linear increase observed in a well-mixed population (**Figure 2A**). Hence even “weak” population subdivision acts to dampen the effects of NFD. In **Figure 2C** we show the approximation in the small migration regime, where the relationship between s and fixation time is reversed.

Note that the effect of frequency dependence on fixation time can essentially be captured by the sign of the derivative of the fixation time at neutrality, s = 0, which can be written as

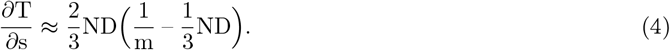

The role of NFD is reversed ( 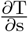 is positive) when NDm ≈ 𝒪(1). Therefore we can define a critical migration rate m^∗^ as

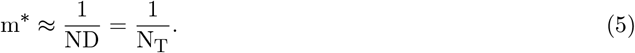

This expression shows that, independent of how the population is divided up into demes, for a given total population size N_T_ = ND, it is the migration rate that critically governs this transition. In **Figure 2D** we show the derivative of the fixation time at neutrality 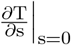 for a populations with a fixed total population of N_T_ = 1000, but with different values for N and D. The analytic approximation given by equation (4) (black solid line) overestimates the critical migration rate when the number of demes is very small (D = 2), since the diffusion approximation we employ assumes a large enough number of demes (D >> 1).

### Non-monotonic dependence of fixation time on deme size

Another prediction that arises from our analytic results is a non-monotonic relationship between the deme size N and the fixation time T. Computing the derivative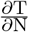,

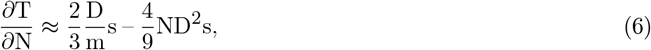

we find that the fixation time is minimized at intermediate deme size 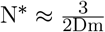.

This expression is similar to that of m^∗^ derived earlier, which means that the non-monotonic pattern occurs in the same regime in which the fixation time is smaller under NFD than neutrality. We show this theoretical prediction of fixation time as a function of deme size under neutrality and NFD in the fragmented (red and blue solid lines, respectively) and the well-mixed limit (grey and orange solid lines respectively) in the insert in **Figure 3**. Specially, the fixation time under NFD is minimized when 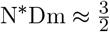 and intersects with the fixation time under neutrality when N^∗^Dm ≈ 3. Beyond this intersection point, NFD promotes coexistence relative to neutrality. Note that under neutrality, theory predicts that the fixation time does not depend on deme size, since migration between demes occurs much slower than fixation within demes. Results from simulations are shown in **Figure 3** and although the theoretical approximations underestimate the fixation time (with increasing N there are increasingly more mutants entering each deme and increasingly stronger effective selection Ns), the prediction of a non-monotonic pattern in the fixation time under NFD remains.

**Figure 3:**
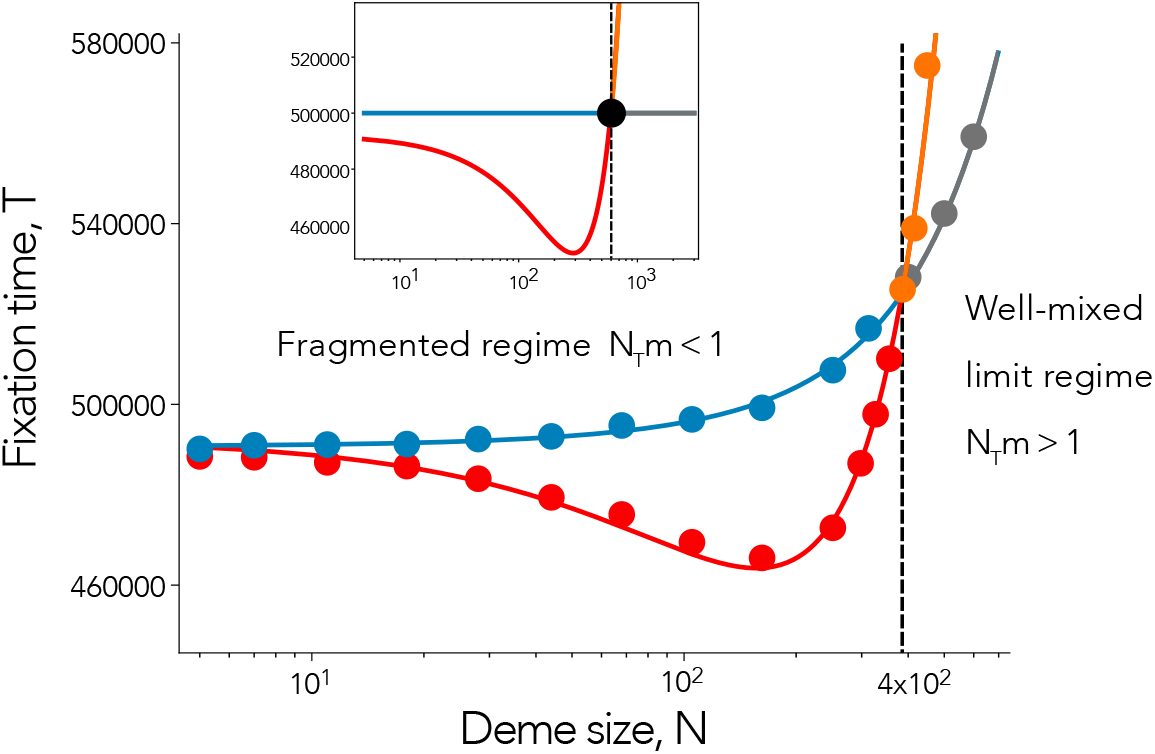
Non-monotonic dependence of fixation time on deme size. Dots represent fixation times estimated from 10^6^ simulation runs, as a function of deme size. We use number of demes D = 50 and migration rate m = 0.0001. Blue points show fixation times under neutrality (s = 0) and red points under NFD (s = –0.001), in the fragmented regime where NDm ≲ 1. Grey dots show fixation times under neutrality (s = 0) and orange points under NFD (s = –0.001) in the well-mixed regime where NDm ≳ 1. Solid lines show simulation results fitted to a quadratic function. The insert shows the corresponding analytic approximations from equation (3), exhibiting similar quantitative behaviours.

### Multispecies coexistence

Thus far we have considered a two variant model and studied average conditional mutant fixation times as a proxy for their coexistence. While this model captures the essence of the interplay between fragmentation and frequency-dependent selection and allows for analytical treatment, it cannot capture other complex dynamics that can occur in fragmented landscapes. A landscape can have demes of different sizes, more complex patterns of connectivity and migration, or multiple species competing in the population.

In this section, to verify that the results from our simple two-variant island model can be extrapolated to more complex ecosystems, we consider a multi-species extension of the model, with demes that can vary in size. We use species richness as the proxy for coexistence, with each species assumed to experience negative frequency-dependent selection as a result of ecological niche differentiation. Since we assume no speciation or migration of new species from an external pool, we analyze the decay of species richness over time, due to loss from random drift.

To construct a meta population with variability in deme size, we utilize a data set of islands that form the Ryukyu islands ecosystem (Matthews et al., 2023). The Ryukyu island system consists of 68 individual islands with areas ranging from 0.041 to 1183 square kilometers. We assume that the area of an island is directly proportional to its population size N and build an island model in which each deme i has size N_i_ = 100A_i_, where A_i_ is the area of island i in square kilometers.

In **Figure 4A** we show results for the well-mixed limit, or “high migration” regime, with N_T_m ≈ 4.5 > 1. Each line shows the decay in species richness over time. In grey, we showcase species richness decay under neutrality for 100 independent runs, with results under NFD shown in orange. In **Figure 4B** we show the histogram of species richness outcomes, after 1e6 generations. In line with expectation, species richness is greater under NFD than neutrality. Meanwhile, in the “low migration” regime (N_T_m ≈ 0.23 < 1), species richness is greater under neutrality compared to the NFD case (**Figures 4c** and **4D**). Hence, the results from our simpler model, and the associated theory, can be extrapolated to more complex ecosystems and different proxies of coexistence.

**Figure 4:**
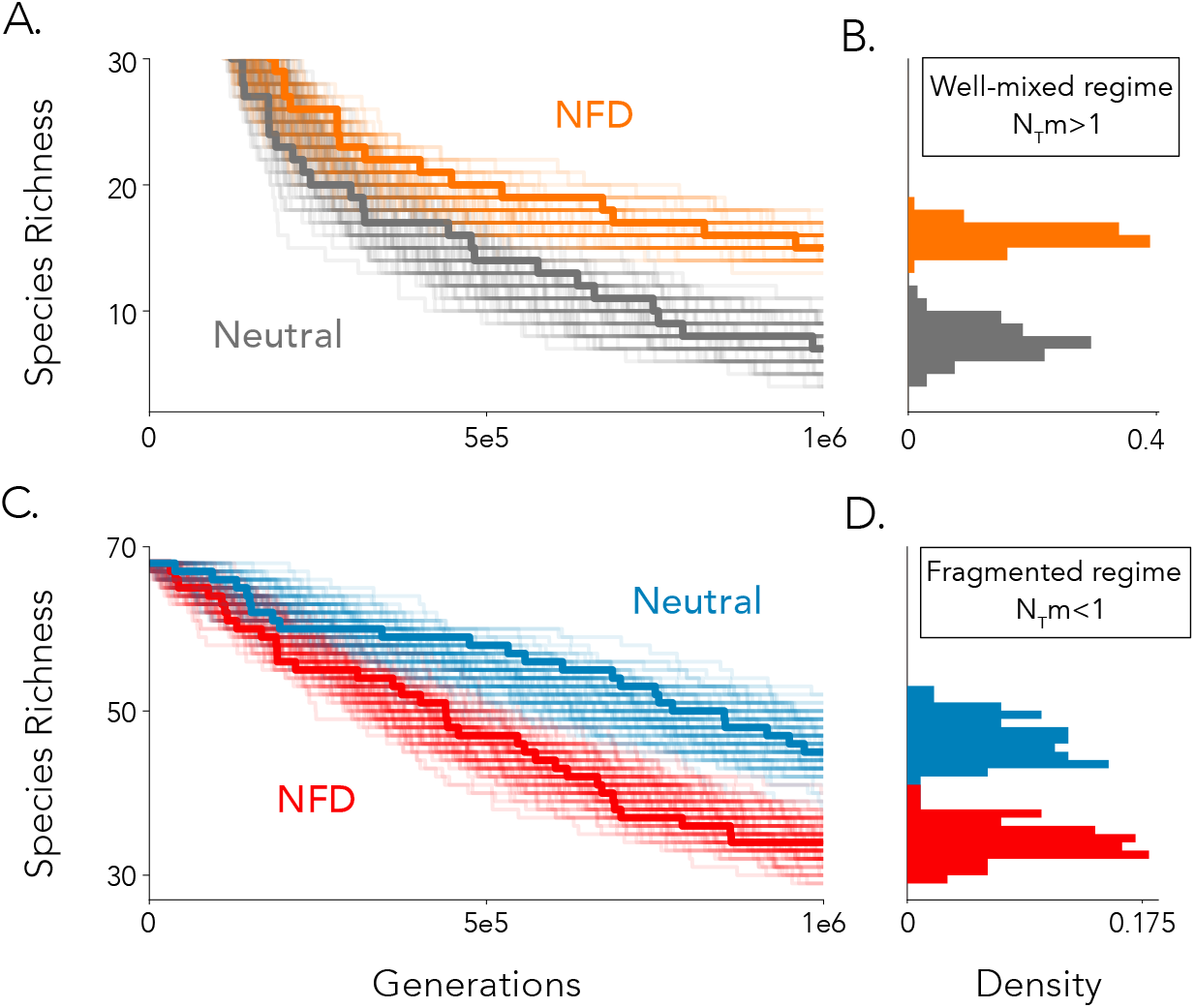
Multispecies model. Each simulation is initialized with a unique species in each deme, such that the initial species richness is 68. **Panel A** shows 100 independent trajectories of the decay in species richness in the high migration regime where m = 1e – 5, NDm ≈ 4.5. Trajectories under neutrality (s = 0) are shown in grey and those under NFD (s = –0.01) in orange. A representative trajectory under both conditions is shown in bold. **Panel B** shows histograms of species richness outcomes after 1e6 generations. **Panel C** similarly shows species richness trajectories in the low migration regime where m = 5e – 7, NDm ≈ 0.23, and **Panel D** the associated species richness outcomes after 1e6 generations.

### Observing negative frequency dependent dynamics in data

We have shown that coexistence in a population increases monotonically with average deme size in three of the studied scenarios: 1) neutrality and high migration, 2) neutrality and low migration, and 3) NFD and high migration. On the other hand, in the case of negative frequency dependent dynamics, coexistence is minimized in fragmented metapopulations with intermediate deme sizes (**Figure 3**). This nonlinear relationship between fixation time and deme size points to a possible test for inferring when coexistence in a metapopulation is maintained by the presence of local negative frequency dependence, over the alternative hypothesis of neutral dynamics.

Using a presence-absence dataset of 99 avian species across the Ryukyu set of islands (data from Matthews et al. (2023), visualization in **Figure 5**), we provide evidence suggesting that avian biodiversity in this fragmented ecosystem is maintained by NFD at a lower level than would be expected under neutrality. Let us first notice that, plotting the species richness values among individual islands, we observe a linear trend with species richness increasing with island area (**Figure 5**).

**Figure 5:**
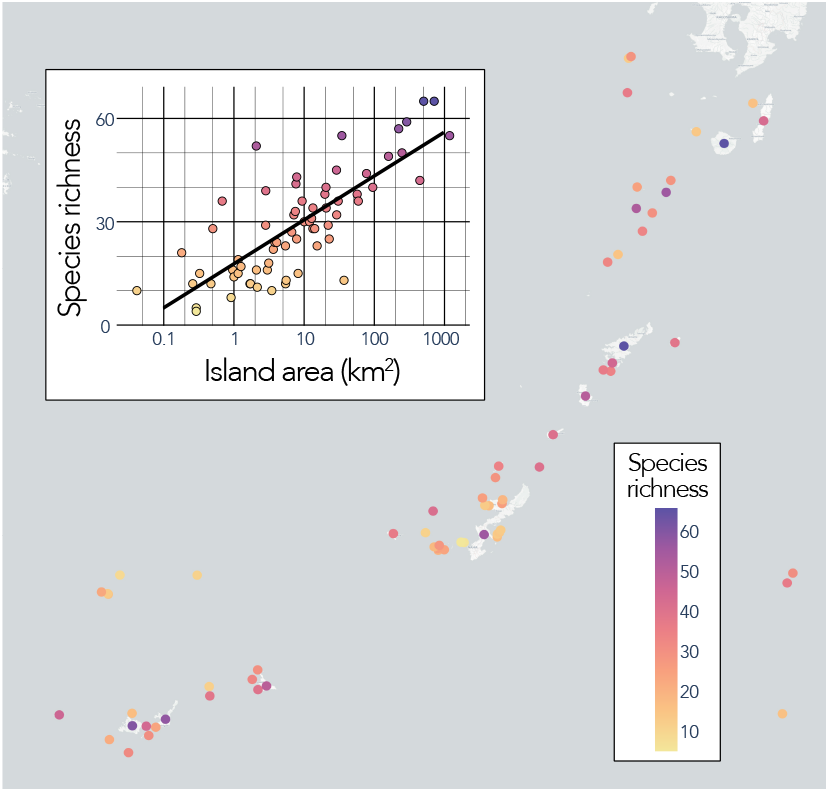
Visualization of data from Matthews et al.(2023). Presence-absence dataset of 99 avian species across the Ryukyu set of islands.

Since in this metapopulation every island (deme) has a different land mass (and population size), direct application of our previous result is not feasible. To understand the relationship between species richness and deme size, let us therefore consider the total species richness in *subsets* of demes of similar size that belong to the larger metapopulation. In order to sample island subsets, we first rank islands by size and then select subsets using a sliding window of size 20. This way of constructing island subsets ensures that we have the broadest range of average island sizes possible. We show that our results are robust to the selected window size of 20 islands in **Supplementary Figure S1**.

In **Figures 6A-C** we plot the total predicted species richness after 1e6 generations, as a function of average deme size, under 1) neutrality and high migration, 2) neutrality and low migration, and 3) NFD and high migration, respectively, and show that species richness indeed increases with deme size, as predicted by our initial simpler model. Here, the solid lines show averages over 100 independent runs of the model, with the shaded areas representing 0.99 confidence intervals. In **Figure 6D** we show results for the case of NFD in the fragmented population regime and, again, observe the non-monotonic relationship between average deme size and species richness. The black dotted lines across the panels indicate species richness in the island subset having the smallest average size in order to highlight that biodiversity initially decreases with increasing average island size only in the regime presented in **Figure 6D**. This theoretical result points to the fact that such a non-linear pattern could be an identifier of a population that is evolving under negative frequency dependent dynamics, given a fragmented population, in the low migration regime.

**Figure 6:**
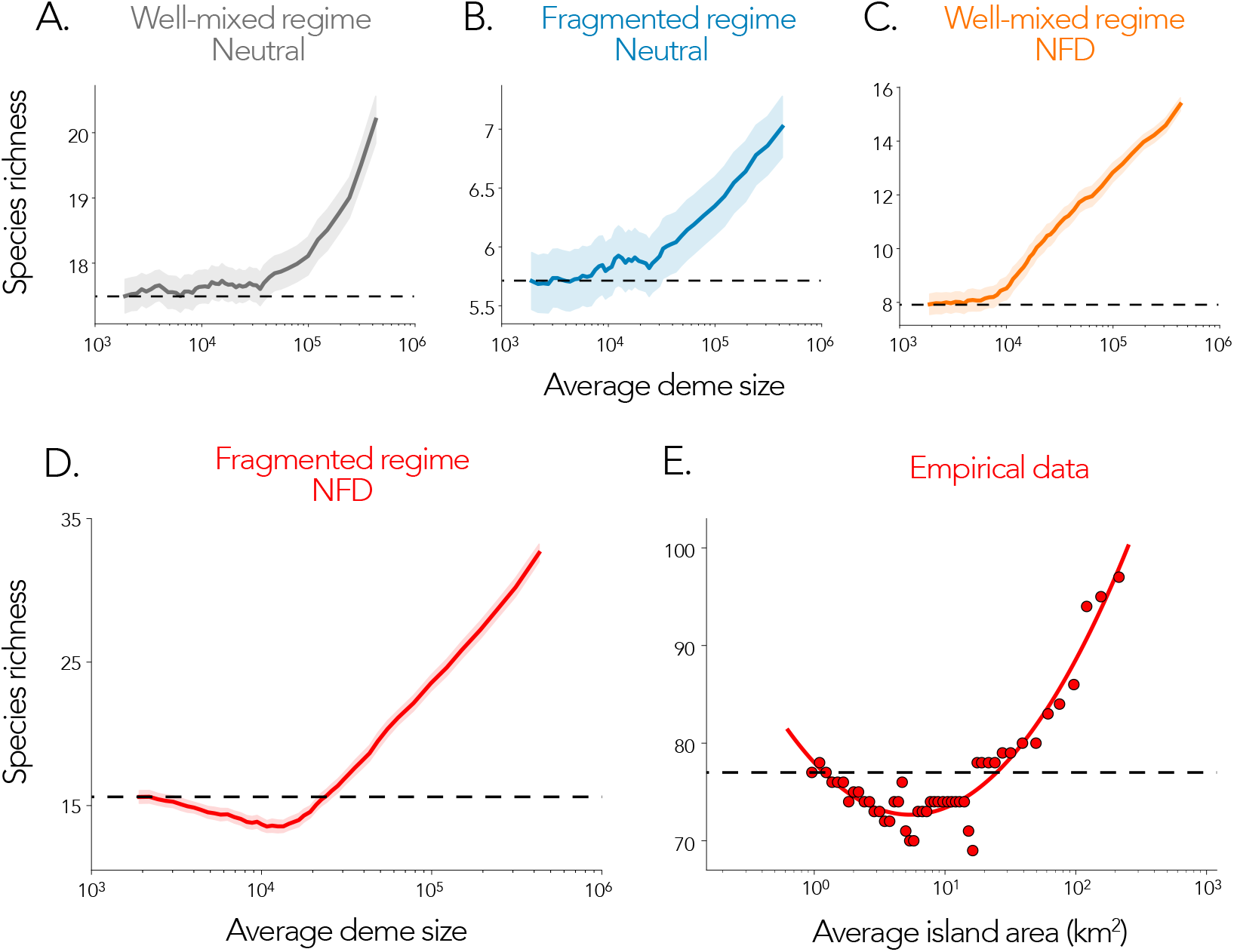
A test for detecting NFD dynamics in fragmented data. In **Figures 5A-D** we show the total species richness using metapopulation deme subsets of size 20 after 1e6 generations, as a function of average deme size under 1) s = 0 and m = 5e – 7, 2) s = 0 and m = 1e – 5, 3) s = –0.01 and m = 1e – 5, and 4) s = –0.01 and m = 5e – 7 respectively. The solid lines show the average over 100 independent runs of the model and the shaded areas represent 0.99 confidence intervals. In **Figures 5E** we show the empirical avian species richness values as a function of average island size using ranked island subsets of size 20. In order to sample fragmented island subsets, we first rank all islands by size and then select subsets of islands of similar size, using a sliding window of size 20 (that is, each fragmented population has exactly 20 islands), and plot the average size in this fragmented island system on the x-axis.

Finally, we perform the same analysis using the true species richness values recorded in the data set of avian species across the Ryukyu islands (Matthews et al., 2023) and observe a similar non-monotonic pattern in species richness as a function of average island area (**Figure 6E**). This suggests that local negative frequency dependent dynamics could be decreasing the species richness of the avian population in these populations, relative to neutrality, since it is only in this regime that such a non-monotonic pattern in species richness is expected. We show that our results are robust to the selected window size of 20 islands in **Supplementary Figure S2**. We also note that the linear trend observed in **Figure 5** highlights that that the nonlinear pattern in species richness is not driven by the individual island level.

## Discussion

Building on the classic island model, we introduce a model of population dynamics driven by frequencydependent selection in fragmented populations. Unlike earlier models of frequency-dependent selection in spatially-structured systems (Luo et al., 2022), our approach considers the effects of drift and demographic stochasticity. In this setting, coexistence is not a binary state but is instead quantified through the average conditional fixation time of a mutant invading a wild-type population (Neuhauser and Pacala, 1999; Murrell, 2010). To understand how spatial fragmentation modulates the effects of frequency-dependent selection, we derive a diffusion approximation for the mutant fixation time. We find that, under sufficient fragmentation and low enough migration, local negative frequency-dependent (NFD) selection can reduce global coexistence, relative to neutrality. This outcome contrasts with results in well-mixed systems, where NFD selection typically enhances biodiversity.

Our results contribute to a growing body of work revisiting the conditions under which NFD selection promotes coexistence. While prior studies have shown that partial niche differentiation can yield lower coexistence than neutrality (D’Andrea and Ostling, 2017), our model goes further: even a fully niche-differentiated community (where each species’ fitness is dependent on its abundance and occupies its own niche) can be outperformed by a neutral system in maintaining diversity, once spatial fragmentation is introduced. Separately, unlike earlier findings in contiguous systems, where coexistence loss arises from interspecific differences in fitness or selection strength (Neuhauser and Pacala, 1999; Murrell, 2010; May et al., 2020; Miranda et al., 2015; Stump and Comita, 2018; Chisholm and Fung, 2020; Smith, 2022; Weiner et al., 2019; Lerch et al., 2023), here, biodiversity loss emerges solely from the interaction of NFD selection with spatial structure, with no variation in selection coefficients (s is fixed for all species). This theoretical result resonates with empirical findings showing that niche-based interactions often fail to predict species coexistence in natural systems (Condit et al., 2012).

This work also aligns with a broader set of ecological and evolutionary models that demonstrate how spatial structure can invert expected outcomes. For instance, positive frequency-dependent selection, typically destabilizing in well-mixed populations, has been shown to prolong coexistence in lattice-structured and patchy landscapes (Molofsky et al., 2001; Molofsky and Bever, 2002). Other processes, sexual selection (M’Gonigle et al., 2012), allee effects (Ferdy and Molofsky, 2002), heterozygote disadvantage (Altrock et al., 2010), and interspecific competition (Ursell, 2021), have similarly shown spatial reversals of their classical effects. Although not the primary focus of this study, our model recapitulates this result. Specifically, we observe that for s > 0, when frequency-dependence is positive, fixation times can exceed neutrality, under low migration.

More broadly, our results offer a new lens on the neutral-versus-niche debate. Specifically, while neutral theory offers a minimal mechanistic model that accurately reproduces key patterns of biodiversity, such as species abundance distributions, it has been criticized for being overly simplistic, since it assumes no niche-based interactions between individuals (Matthews and Whittaker, 2014), despite evidence that such interactions are widespread (Purves and Turnbull, 2010). Our analysis reveals a possible explanation: in fragmented systems, populations subject to NFD selection can appear effectively neutral at the global scale. Neutral theory has also been criticized due to its inability to reproduce certain other key characteristics of natural populations. For instance, neutral theory significantly overestimates fixation times and underestimates the magnitude of abundance fluctuations (Kalyuzhny et al., 2015). Our model of fragmented populations also offers a solution to these limitations, since it predicts that fixation times may be shorter and the variance in per-generation change in the mutant frequency can be larger under NFD selection, compared to neutrality.

We also extend our model to a multispecies regime and confirm that our findings hold when coexistence is measured via species richness. Specifically, using a metapopulation model based on 68 islands from the Ryukyu archipelago, we find that under low migration, species richness is lower under NFD selection than under neutrality, mirroring predictions from the two-species model. To test these ideas empirically, we derive a signature of the low-migration NFD regime, by building on the classic species-area relationship, which demonstrates that species richness within islands generally increases monotonically with the area of the island (Preston, 1962). While species richness typically increases with island area (Preston, 1962), we find that under NFD selection with limited migration, total richness follows a non-monotonic trend: it is minimized at intermediate total area. This contrasts with the monotonically increasing pattern seen in all other regimes. We apply this test to a presence–absence dataset of 99 avian species across the Ryukyu Islands (Matthews et al., 2023) and observe the predicted non-monotonic relationship, suggesting that this ecosystem is governed by NFD selection.

The simplicity of the models we study here constitute both a strength and a weakness. The models are simple enough to provide a unifying proof of concept for the resulting evolutionary dynamics, but they also omit several ecological complexities of real fragmented metapopulations. One such limitation lies in the assumption of uniform migration rates between all the demes in the metapopulation. More realistically, migration rates between islands can be variable and moreover, the islands themselves need not all be connected and can instead form a network of connected demes, with varying topological properties, shaped by factors such as distance and matrix quality (Gallé et al., 2022). We also assume no interspecific variation in selection strength (s) and no intrinsic fitness differences, both of which are known to impact coexistence outcomes (Stump and Comita, 2018). All of these modeling extensions are necessary for a unifying theory on the role of negative frequency dependence in shaping patterns of coexistence and future work will need to explore their roles in shaping evolutionary dynamics in fragmented ecosystems.

## Model and Methods

### Frequency-dependent dynamics in a fragmented population: two variant model

To a model a spatially-structured population of species competing under frequency-dependent selection, we consider an extension of Wright’s classic island model and assume a population wild-type and mutant variants that are structured into D demes of N individuals each linked by migration occurring with probability m. This means each that generation the offspring’s parents come from within the deme with probability (1 – m) and from the overall population with probability m. The population composed of two different types of individuals: wild-type and mutant variants, with further mutation neglected.

Following Molofsky and Bever (2002), we assume that reproduction within demes is biased by local frequency-dependent selection, such that the mutant fitness in deme i is given by

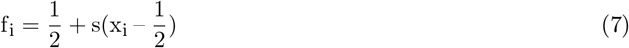

where x_i_ is the mutant frequency in deme i and s is a frequency-dependent selection coefficient. The wild-type fitness in deme i follows follows the same form, where we replace x_i_ with (1 – x_i_) and get 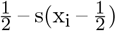, which allows us to write the relative mutant frequency 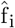 as

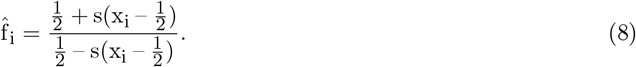

For each deme, to create the next generation, individuals are sampled from within the deme with probability (1 – m) and from the entire population with probability m, followed by local frequency dependent selection. Therefore, under neutrality, an individual in deme i is of the mutant type with a probability given by

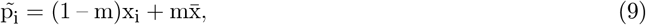

where x_i_ is the mutant frequency in deme i and 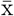 is the mean population-wide mutant frequency in the previous generation. When the sampling of alleles is further biased by local frequency-dependent selection according to equation (7), the mutant sampling probability becomes

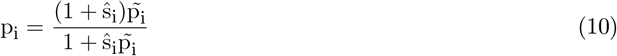

where 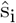 is the relative mutant selection coefficient (Cherry and Wakeley, 2003). To determine 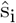 for our model, using a Taylor expansion around s = 0, we can write the relative fitness of the mutant allele in deme i as

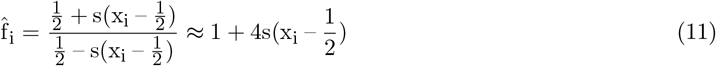

and therefore 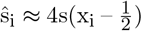.

### Frequency-dependent dynamics in a fragmented population based on the Ryukyu island ecosystem: multispecies model

To verify that our results can be extrapolated to more complex ecosystems, we consider a multispecies extension of the two-variant island model. Specifically, we relax two assumptions: 1) there may be more than two species and 2) deme sizes may vary.

Let us assume that the fitness of species *σ* in deme i is given by

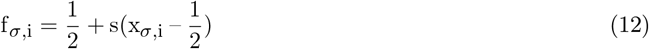

where x_*σ*,i_ is the frequency of the species in deme i, similar to equation (7). Under this fitness function, the equilibrium state of a well-mixed population still occurs when all the species have equal frequencies because this is the condition under which they all attain the same value of f_*σ*,i_. Reproduction and migration are implemented in the same as the two variant model above. In this multispecies model, we focus on species richness as the main proxy for coexistence. Since we assume no speciation events that can create new species or migration of new species from an external pool, the species richness will only decay over time, because of loss of species from random drift. We initialize the population with a unique species in each island and analyze this decay of species richness over time.

In order to understand the maintenance of biodiversity in an island ecosystem with varying island sizes, we model the metapopulation after the structure of the Ryukyu island ecosystem. The Ryukyu islands are made up of 68 individual islands that vary in area ranging from 0.041 square kilometers to 1183 square kilometers. We make the assumption that the area of an island is directly proportional to the population size on that island, N. Specifically, we represent the Ryukyu island population using an island model in which each deme i has size N_i_ = 100 · A_i_, where A_i_ is the area of island i in square kilometers. We select a scaling factor of 100 for N_i_, because this is the minimum value at which all islands have a size that is greater than 1.

## Data and code availability

Custom scripts were used for the simulations and data analyses. All C++ and Python simulation code is available on Github at XXX (repository to be made public upon acceptance and publication of the manuscript). All packages used for analysis and visualization are open-source.

## Acknowledgments

This research was done using resources provided by the Open Science Grid, which is supported by the National Science Foundation award 1148698, and the U.S. Department of Energy’s Office of Science.

## Supplementary Appendices and Figures

### Supplementary Appendix 1. Conditional fixation time in an island model with frequency-dependent selection

Here we derive analytic approximations for the conditional mutant fixation time, in the limit of weak migration and selection. Assuming m << 1 and s << 1, we write the mean per-generation change in the mutant frequency in deme i as

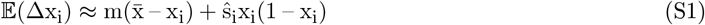

and, subsequently, the population wide mutant frequency, 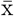, as

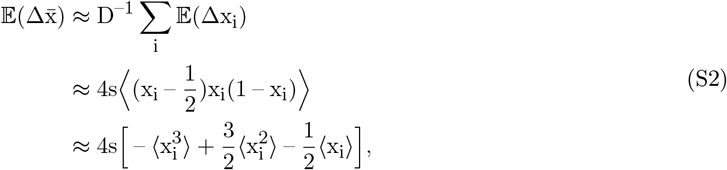

where we assume that the expected change due to the migration component of 𝔼(Δx_i_) sums to 0 across demes, i.e. 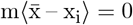 (Cherry and Wakeley, 2003).

In order to express 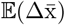 in terms of 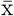, we assume that each individual deme behaves as an island population, coupled to a much larger continent population via migration, such that the population-wide mutant frequency 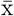 is at equilibrium on the timescale of changes in the local mutant frequencies x_i_. We must therefore first write the distribution of allele frequencies in an island-continent model with frequencydependent selection. Consider a single island population that receives migrants from a much larger continent population at rate m. Each generation, the island population undergoes a migration step followed by a frequency-dependent selection step. We can approximate the mean per-generation change in the mutant frequency as

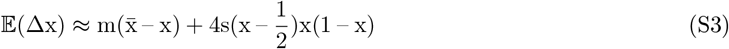

(Cherry and Wakeley, 2003) and the variance as

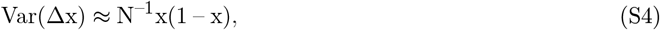

where × is the mutant frequency in the island, N is the island size, and s is the strength of frequencydependent selection. We assume that the continent mutant frequency 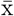 changes much more slowly than the island mutant frequency. We may then write the mutant frequency distribution as

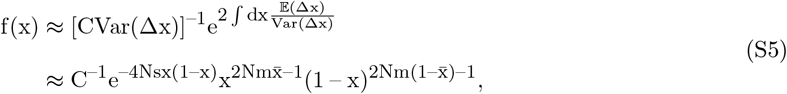

where C is a normalization constant (see, for example, equation (4.45) in Ewens (2004)). Note that we have absorbed a constant factor of N into C. To enable tractable computation of C, we can first rewrite f(x) as

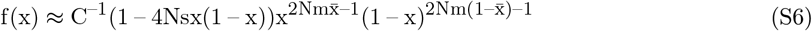

using a Taylor expansion around s = 0, up to 𝒪(s) terms. We can then compute C as

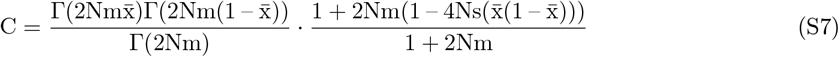

using the normalization condition 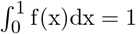, where Γ(·) is the gamma function.

After deriving this island-continent mutant frequency approximation, we consider x_i_ approximately distributed according to (S6), allowing us to write 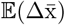 using the associated moments. We write the first three moments of (S6) as

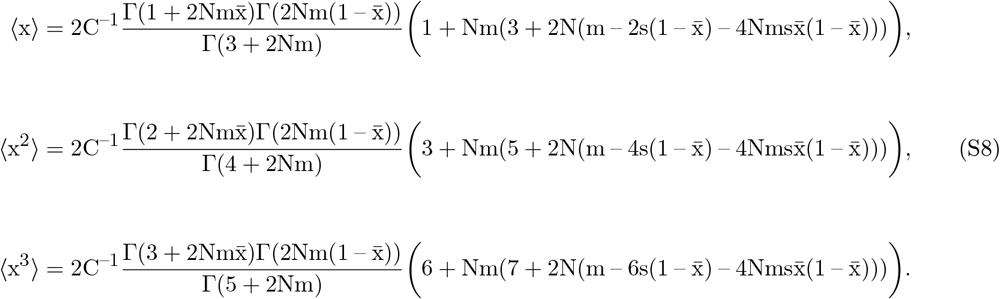

After plugging in C from equation ((S7)) and a Taylor expansion around s = 0 up to 𝒪 (s) terms, and m = 0 up to 𝒪 (m^2^) terms, we can write

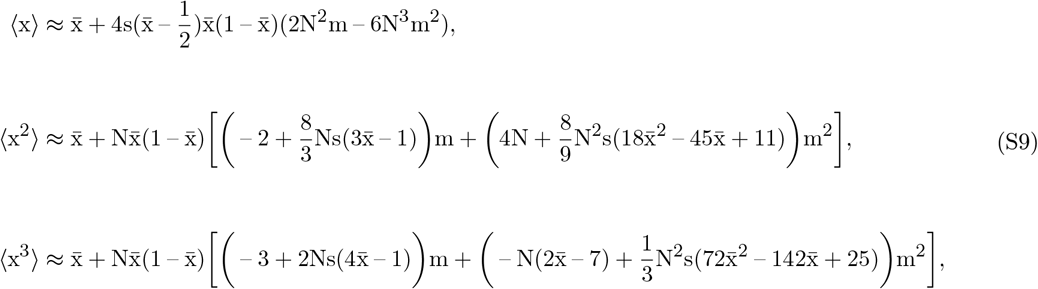

which allows us to write 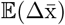 as

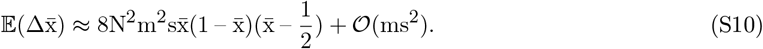

Similarly, we can write the variance of the per-generation change in the mutant frequency in deme i as Var(Δx_i_) ≈ N^−1^p_i_(1 – p_i_), and that of the population-wide mutant frequency as

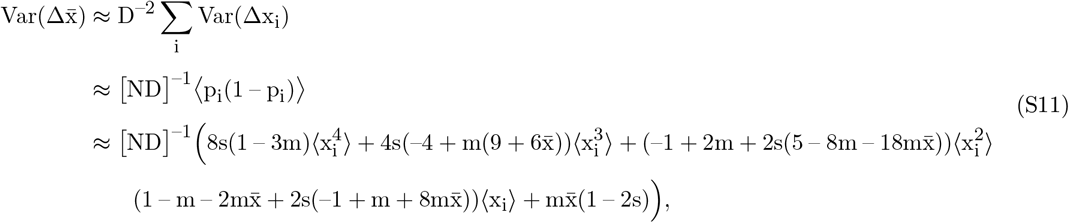

where, in the last step, we again use a Taylor expansion around s = 0 and m = 0 and retain only up to 𝒪 (ms) terms. After computing the fourth moment of (S6) in a similar way,

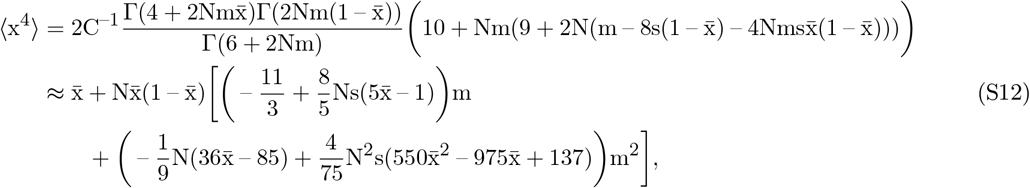

we can write 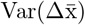 as

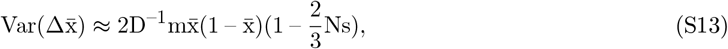

where we retain only up to 𝒪 (ms) terms. We can now write the Kolmogorov backward equation as

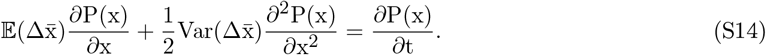

The fixation probability of the mutant type at an initial frequency x, P(x), satisfies the Kolmogorov backward equation at steady state when ∂P(x)/∂t = 0. We write the solution to this equation as

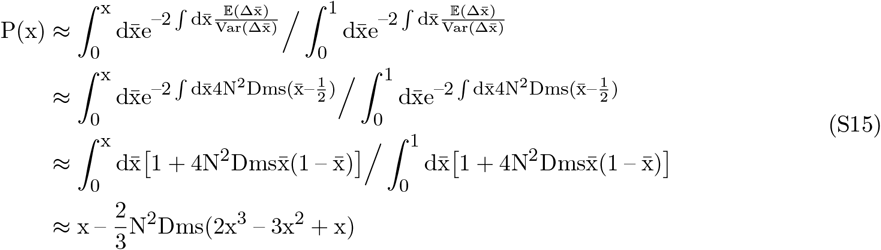

using a Taylor expansion around s = 0 and m = 0 up to 𝒪 (ms) (see, for example, equation (4.17) in Ewens (2004)). Similarly, the mean fixation time of the mutant type, given an initial frequency x, T(x) satisfies

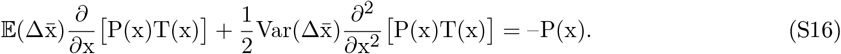

The solution to this equation may be written as

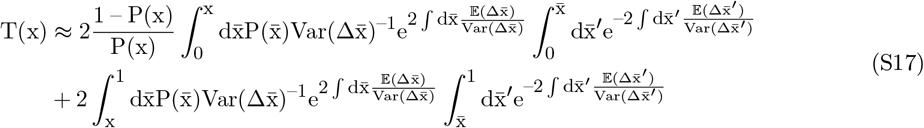

(see, for example, equations (4.49) and (4.50) in Ewens (2004)). However, since we assume the population starts with a single mutant individual x = 1/ND, we may ignore the contribution arising from 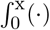 for large ND, since x → 0. This allows us to write

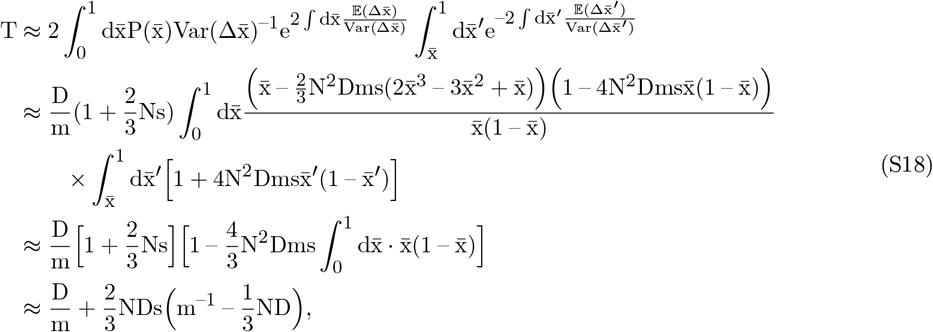

again, using a Taylor expansion around s = 0 and m = 0 up to 𝒪 (ms) terms.

## Supplementary Figures

**Supplementary Figure S1:**
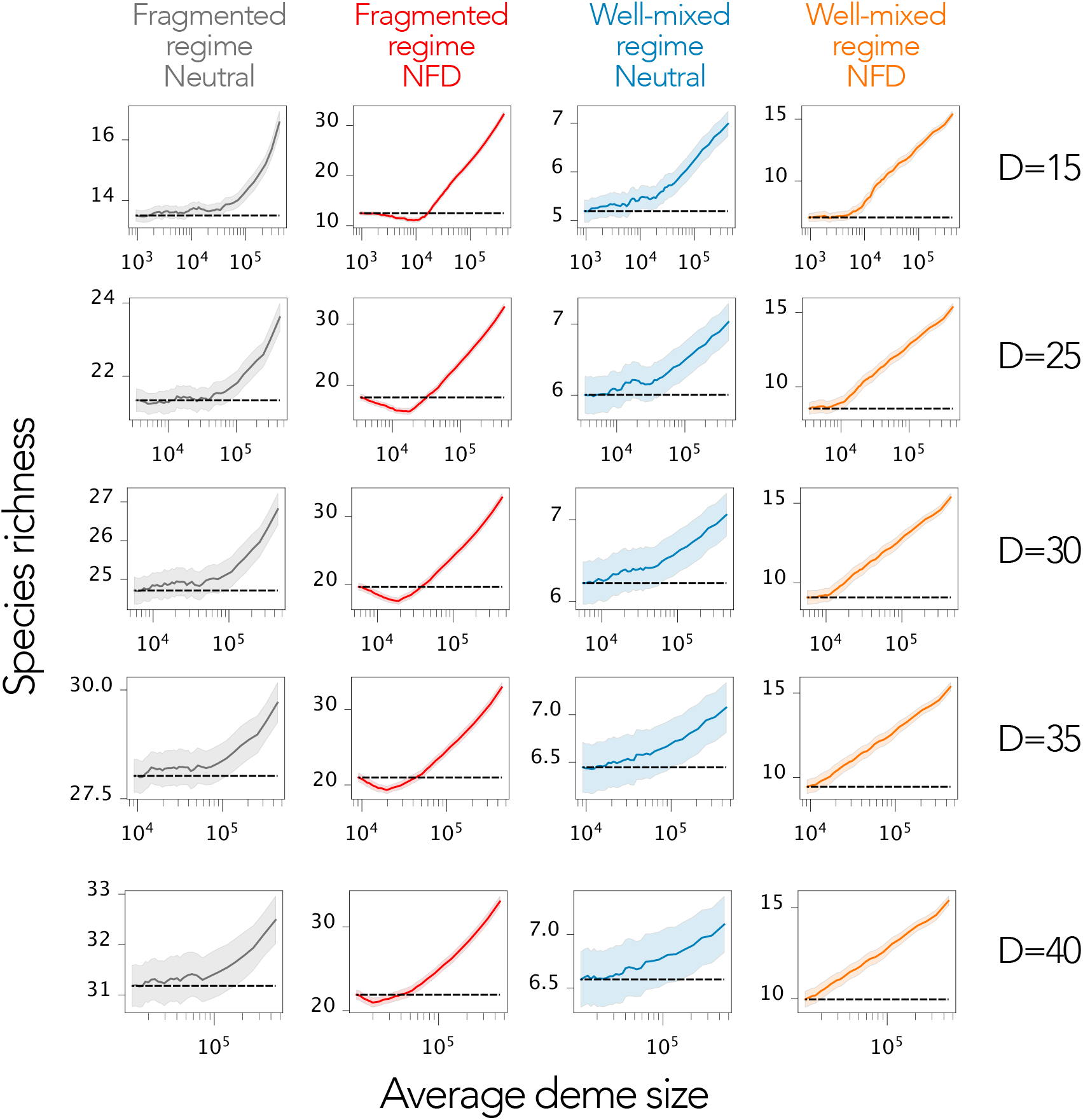
Robustness to ranked island window size, simulations. Same parameters as **Figure 6**. In order to sample fragmented deme subsets, we first rank demes by size and then select subsets of demes of similar size, using a sliding window of size D (with D as in the panels, ranging from 15 to 40) and plot the average size of the demes in the subset on the x-axis. The nonlinear pattern is present regardless of sample size.

**Supplementary Figure S2:**
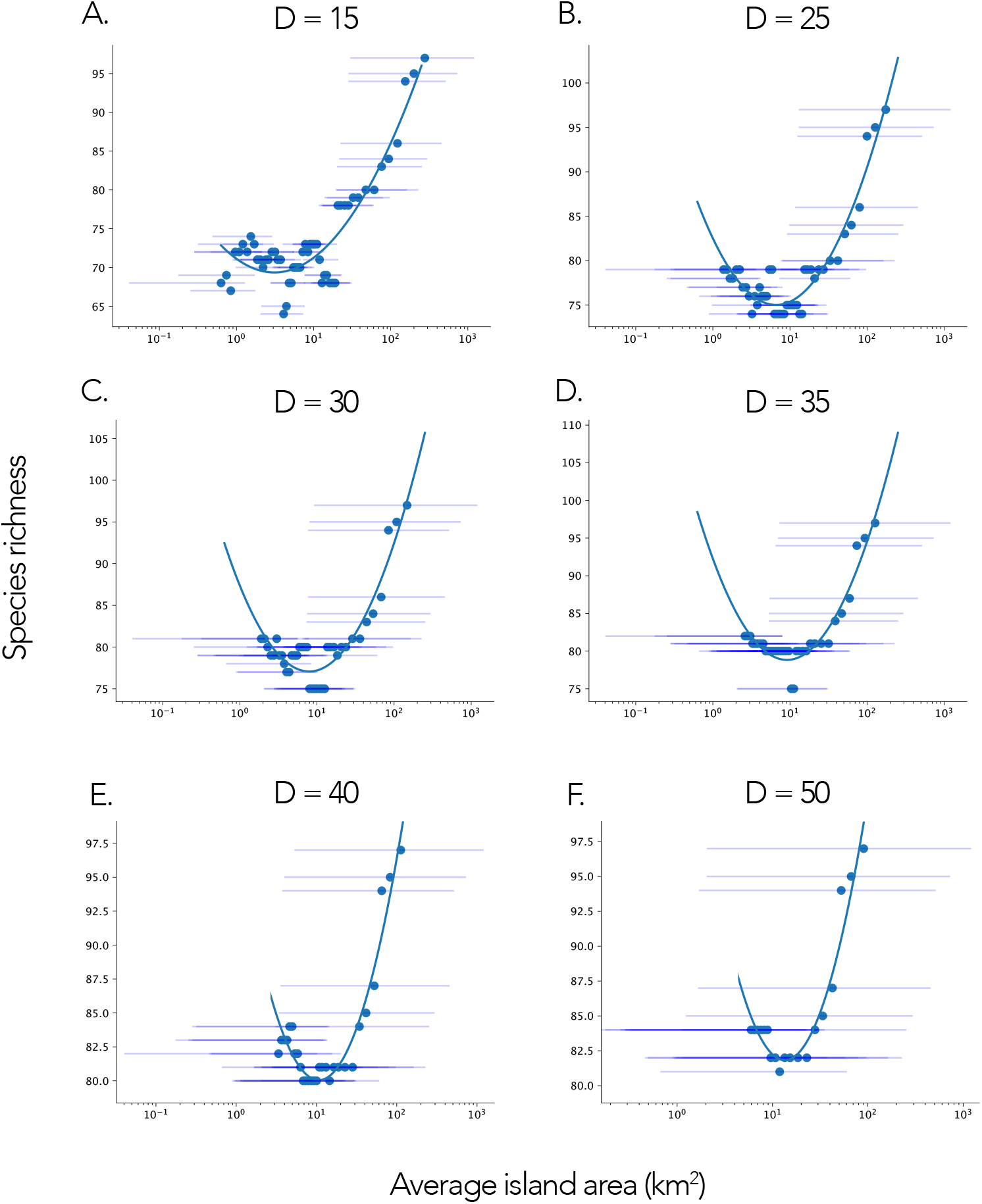
Robustness to ranked island window size, empirical data. Same parameters as **Figure 6**. In order to sample fragmented island subsets, we first rank islands by size and then select subsets of islands of similar size, using a sliding window of size D (with D as in the panels, ranging from 15 to 50) and plot their average subset size on the x-axis. The nonlinear pattern is present regardless of sample size.

